# Differential warming at crown scale impact walnut primary growth onset and secondary growth rate

**DOI:** 10.1101/2024.03.25.586536

**Authors:** Nicolas Dusart, Bruno Moulia, Marc Saudreau, Christophe Serre, Guillaume Charrier, Félix P. Hartmann

## Abstract

Trees are exposed to significant spatio-temporal thermal variations, which can induce intracrown discrepancies in the onset and dynamics of primary and secondary growth. In recent decades, an increase in late winter and early spring temperatures has been observed, potentially accelerating bud break, cambial activation and their potential coordination. Intracrown temperature heterogeneities could lead to asymmetric tree shapes unless there is a compensatory mechanism at the crown level.

An original warming experiment was conducted on young Juglans regia trees in a greenhouse. From February to August, the average temperature difference during the day between warmed and control parts was 4°C. The warming treatment advanced the date of budbreak significantly, by up to 14 days. Warming did not alter secondary growth resumption but increased growth rates, leading to higher xylem cell production (twice as many) and to an increase in radial increment (+80% compared to control). Meristems resumptions were asynchronous without coordination in response to temperature. Buds on warmed branches began to swell two weeks prior to cambial division, which was one week earlier than on control branches. A difference in carbon and water remobilisation at the end of bud ecodormancy was noted under warming. Overall, our results argue for a lack of compensatory mechanisms at the crown scale, which may lead to significant changes in tree architecture in response to intra-crown temperature heterogeneities.

Highlight: When tree are submitted to asymmetrical warming, it leads to early budbreak and enhanced cambial activity for warmed branches

## Introduction

Temperature is a major environmental driver of plant development, to such an extent that reverse phenological models are able to reconstruct past temperatures from historical grape ripening data (Chuine *et al*., 2004). In the last decades, changes in temperature regimes are observed (Calvin *et al*., 2023). An increase in temperature in autumn could delay endodormancy induction (Beil *et al*., 2021; Charrier, 2022); in late winter and early spring, it could hasten the onset of primary and secondary growth *i.e.* budbreak (Chuine, 2010), cambial activation (Begum *et al*., 2013), and their coordination (Savage and Chuine, 2021 for review).

Mechanisms regulating budbreak are well described and their regulation is characterised by different stages : paradormancy, endodormancy and ecodormancy (Lang, 1987). Paradormancy is mostly determined by apical dominance, which dependency on temperature has remained unclear. Endodormancy is induced by cold night temperature and low photoperiod and released by chilling exposure. Once endodormancy is over, ecodormancy is released by warm temperatures and, eventually, long photoperiod. Differences in chilling and forcing requirements to release endo- and ecodormancy, respectively, explain inter- and intraspecies variability in blooming and budbreak dates (Chuine, 2010; Charrier *et al*., 2011). The resumption of cambial activity is likely to be affected by a sequential exposure to chilling and forcing temperatures, as formalised in a modelling approach in northern hemisphere conifers (Delpierre *et al*., 2019). Nevertheless, it had been suggested that cambial rest is deeper and longer in deciduous than evergreen (Oribe and Kubo, 1997; Begum *et al*., 2007; Kudo *et al*., 2014). For poplar, a period of several days above a threshold of 15°C daily maximum temperature before cambial reactivation had been proposed (Begum et al., 2008). Beyond these differences in phenological control, it is obvious that the response of growth resumption to temperature differs between primary (buds) and secondary (cambium) meristems. In buds, the integration of temperature along a significant period is appropriate in predicting the budbreak date. In cambium, on the contrary, the date where temperature exceeds a threshold is more relevant to predict the date of resumption of secondary growth (Antonucci *et al*., 2015).

A further complexity in the analysis of temperature responses in primary and secondary meristems lies into the various time sequences between primary and secondary growth resumptions across species and functional types. Three different patterns are observed with respect to wood anatomy and are genetically controlled: (i) cambium reactivation before budbreak, in many vesselless evergreen species, like *Picea Abies* (Kraus *et al*., 2016), or in ring-porous deciduous (Gričar *et al*., 2020); (ii) synchronicity, *e.g*. in *Abies balsamea* or *Picea mariana* (Antonucci *et al*., 2015); (iii) budbreak before stem increment, in diffuse-porous trees like *Fagus sylvatica* (Čufar *et al*., 2008; Kraus *et al*., 2016) or *Populus tomentosa* (Li *et al*., 2013). A functional hypothesis has been proposed connecting the timing of growth resumption with the capacity to transport carbon and water for the sake of resuming primary growth; indeed, the species that are unable to repair winter embolism (eg, ring-porous *Quercus*, Tyree and Cochard 1996) must resume cambium growth before budbreak (Savage and Chuine, 2021). Coordination between meristem is still unclear as disbudding experiment revealed that cambial reactivation and vessel differentiation can occur independently of budbreak (Kudo *et al*., 2014).

Temperature fluctuations influence cellular biological activity (Peterson *et al*., 2007). Increasing temperature can enhance biological processes such as respiration, enzymatic activity, and gene expression rates up to an optimum threshold, beyond which further rises can lead to their decline(Sharpe and DeMichele, 1977; Zwieniecki *et al*., 2015). All of these processes depend on an energy supply. For deciduous trees, carbon reserve constituted before the leaves fall in the form of non-structural carbohydrates (NSC—soluble carbohydrates and starch) may be among the key mechanisms regulating plant internal dormancy clocks (Tixier *et al*., 2017*a*; Amico Roxas *et al*., 2021). Indeed, management of these NSC reserves during winter dormancy should assure an adequate supply for spring growth resumption. Besides, the onset of active primary or secondary growth is acting as strong sinks for the carbohydrate in a time where only reserves can act as a source (Charrier *et al*., 2018). As the activity of amylase, for example, is very temperature dependant (Zolfaghari *et al*., 2005), the supply of soluble sugars may be affected by temperature, which could drive the response of growth to temperature. This suggests that the level of carbohydrates reserves and associate metabolism assuring balance between active (hexose) and storage (starch) forms may be involved in developmental stage change and therefore in the response to temperature of primary and secondary growth. However, recent researches have emphasized that the influence of sugars on meristem activation may act through signalling, that would anticipate trophic regulation. This is well-documented for bud break in *Rosa hybrid*, *Prunus persica*, and *Juglans nigra* (Marquat *et al*., 1999; Bonhomme *et al*., 2010; Bertheloot *et al*., 2020).

The temperature of organs and tissues can also be spatially heterogeneous at tree scale. Indeed, trees are facing temperature heterogeneity because of the asymmetry of solar radiation. The face of the tree (and of each organ) that is more exposed to solar radiation has a higher radiative budget than the opposite one (Saudreau *et al*., 2011; Peaucelle *et al*., 2022), leading to important thermal differences at the tree scale. These differences depend on the organ size and the microenvironment. Among all organs, the trunk exhibits the greatest difference between its North and South sides, ranging from 10 to 20 °C depending on the diameter (Sakai, 1966; Sheppard *et al*., 2016; Musselman and Pomeroy, 2017). The highest differences between North and South sides occur on days with strong insolation and low wind. This range of variation involves freezing temperature typically leading to sunscald (Yang *et al*., 2020). As smallest structures, buds were assumed to be in equilibrium with the air, although non-negligible (up to 6°C) local temperature variations have been measured (Grace, 2006; Peaucelle *et al*., 2022). Temperature differences across the crown have been observed during summer and autumn but not in spring (Zahnd *et al*., 2023), leading to homogeneous bud breaking, but less likely to affect branch primary elongation (Schiestl-Aalto *et al*., 2013).

Finally, would heterogeneous intra-crown temperature lead to heterogeneous bud outgrowth, cambium reactivation, and primary and secondary growth rate and duration, resulting in asymmetric time course and shape at the tree level or do the integration of such local responses compensated at the crown level? In this study, we therefore investigated jointly the responses of buds and cambium to temperature and to its time fluctuations and spatial heterogeneity, from bud ecodormancy to the end of vegetative season. More precisely, we assessed five hypotheses:

1. The resumption of growth for primary and secondary meristems respond independently of each other to temperature.
2. Primary and secondary growth rates increase similarly with temperature.
3. The cessation of growth is independent of temperature and is not coordinated between the meristems.
4. Spatial temperature gradients across the crown result in asymmetric growth.
5. Sugars released from reserves in the stem can explain the changes in primary and secondary growth kinetics.

We present an original set-up of warming experiment on *Juglans regia* young trees in greenhouse to test the effect of temperature asymmetry. Determining primary and secondary growth resumption, rate and cessation together and accurately was a methodological challenge. Bud phenology was determined by two different methods, automated with timelapse camera or visual observation. Secondary growth was assessed continuously with an automated dendrometer and at specific time points by cytological analysis. For both meristems, the developmental stage was precisely defined to characterise growth resumption and cessation. At a more physiological level, carbohydrates and water content in different parts of the branch were also measured to explore carbon and water balance during the whole experiment.

## Material & Methods

### Plant material and thermal treatment

Forty five-year-old walnut trees (*Juglans regia* cv Franquette) were grown in 80 L containers with a local soil mixture (1:3 black peat, 2:3 Terre de Limagne, volume). Trees were grown under natural conditions until the end of December so that chilling requirements were met and endodormancy released. In the first week of January, the trees were transferred to a greenhouse in which air temperature was maintained above 5 °C. Light irradiance was monitored using three Photosynthetically Photon Flux Density (PPFD) sensors (PAR/CBE 80, Solems, Palaiseau, France, Supplementary Fig. S1).

For each tree, the southern face of the trunk and all the branches were covered using aluminium tape (425,3M, Saint Paul, Minnesota, USA). On branches, tape was wrapped in a spiral around the branches, leaving *ca.* 1 cm of uncovered bark between spires. Heating resistances, made of constantan wires (diameter 0.2mm), were mounted above the aluminium tape and fixed by a second layer of aluminium tape on the top of the first one (Fig. 1 A&B). Electrical current was supplied to the constantan wire so that the heat was conducted by the tape to half the trunk and half of the branches. Branches were grouped according to their volume (length, diameter) to modulate the tension of the electrical supply so a similar warming was applied. Across groups of branches, the applied tension ranged from 8 to 14 V generated by a 60 W universal regulated modular power supply (ALE2902M, elc, Annecy, France). To avoid excessive warming during sunny days and overnight, a controller composed of a light sensor connected to an Arduino board (UNO rev 3, Arduino, Scarmagno, Italy) switched on the power supply. The warming was applied only when PPFD was between 50 and 800 µmol.m^-2^.s^-1^. The temperature of two branches per tree (warmed and control) was monitored using type-T thermocouple connected to a datalogger (CR1000, Campbell, Logan, Utah, USA) mounted in contact with the bark of the last growth unit (Fig.1 E).

**Figure 1:**
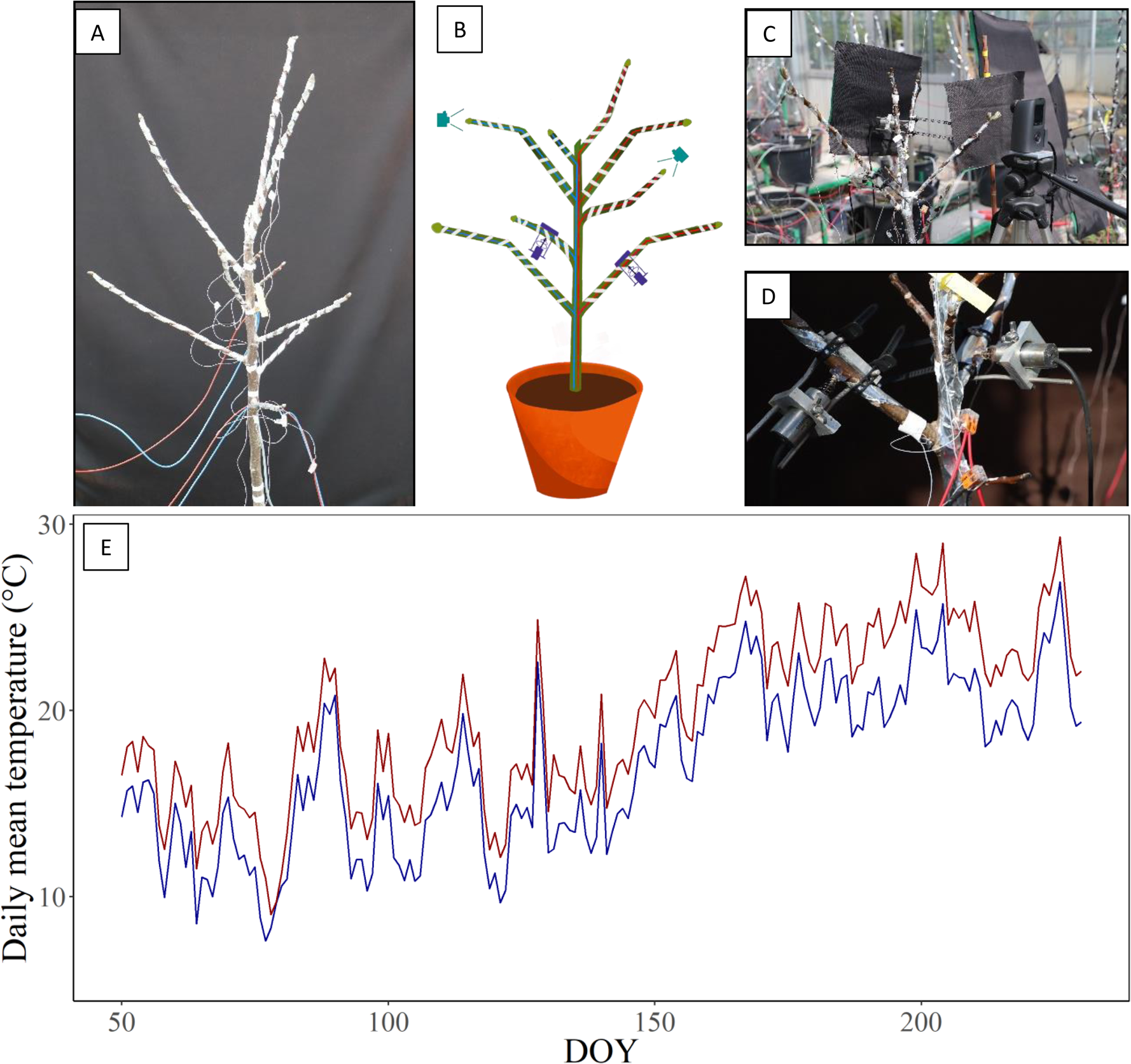
Photography of the warming system (A) schematic view of a fully experiment tree (B), timelapse system (C) and automatic dendrometer (D). Fluctuation of daily mean temperature during the experiment with control branches in blue and warmed branches in red (E).

### Budbreak and primary growth

The phenological stages of buds were monitored on five trees using timelapse cameras TLC200 Pro (Brinno, Netherlands) (Fig. 1B&C). A picture was taken every 2 hours during daytime, and the brightest one per day was selected for image analysis using Fiji software (Schindelin *et al*., 2012). The area of each bud was measured as the number of pixel per bud, extracted by manually modulating the contrast and the brightness of the selected image. The date when the area of the bud increased by 50 % was defined as an index for the onset of budbreak.

During the same period, phenology of each terminal bud was visually observed every 2 days according to BBCH scale (Meier, 2018). With a focus on the early stages from stage 00, dormant bud, to stage 10, first leaf emergence, including stage 07 corresponding to budbreak and stage 09 when the first leaves become detached. After buds flushed (stage 10 BBCH), the length of the growing unit was measured every two weeks on ten pairs of branches (warmed and control).

### Secondary growth

From February to August, high-resolution dendrometer DF 2.5 LVDT (Solartron Metrology, Bognor Regis, United Kingdom) monitored the variation in diameter of ten pairs of branches (warmed and control) (Fig. 1B&D). Sensors were connected to a datalogger (CR3000, Campbell, Logan, Utah, USA). Data were recorded every 30 minutes.

### Samplings and physiological analysis

Every two weeks, branches were sampled for water content, carbohydrates and cytological analysis of five trees. The branches were divided as follows (Fig. 2): 1) Before budbreak, all the buds of the branch (3 buds average) were immediately excised from the stem and pooled. After budbreak, when the new growth unit (n) reached 10 cm long, a segment of 10 cm from the apex was sampled (leaves removed) for analysis. 2) the 2020 growth unit (n-1) were sampled for cytological analysis and as younger parts were too small for both analyses, a 10 cm segment of the 2019 growth unit (n-2) was sampled for carbohydrates content.

**Figure 2:**
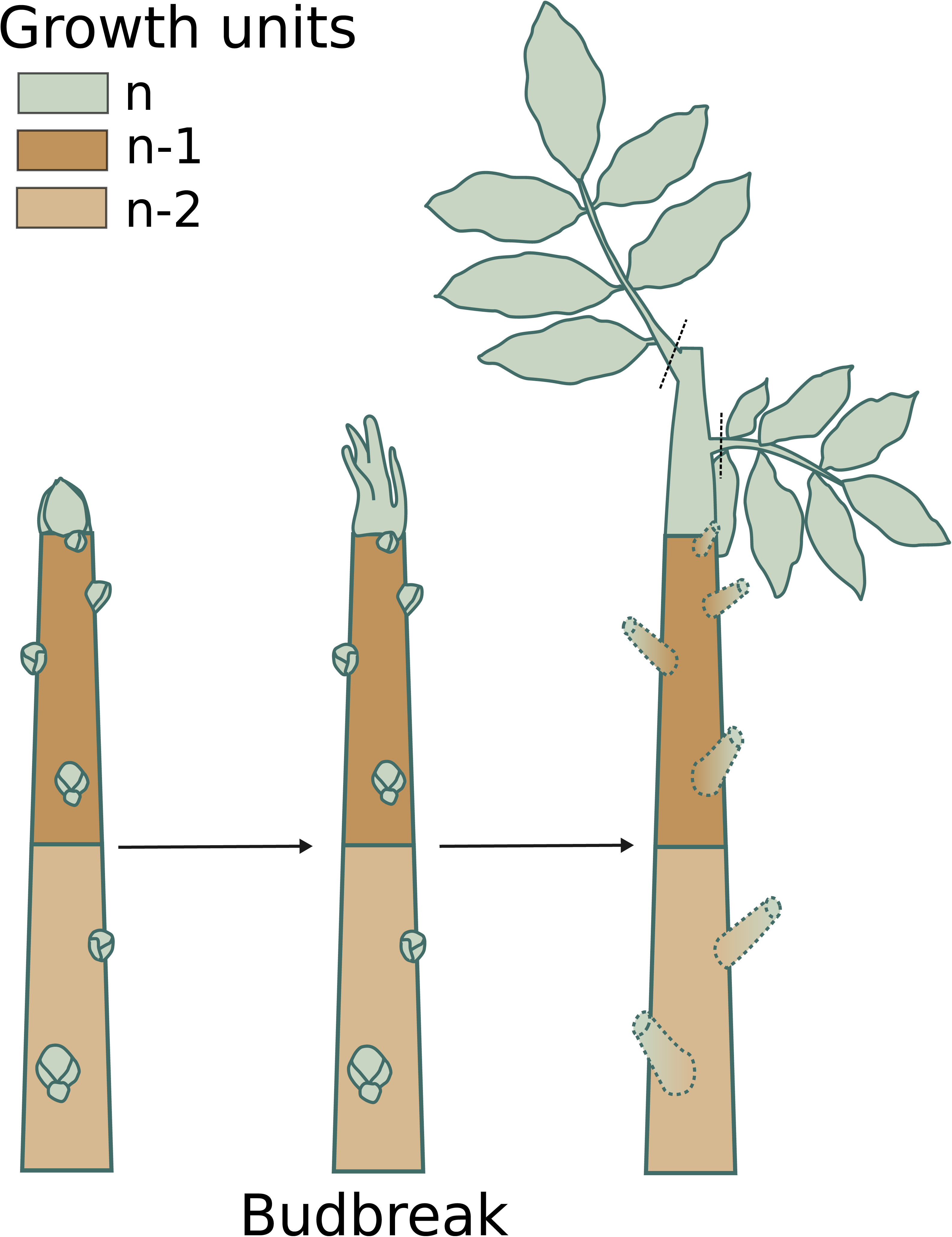
Schematic representation of sampled growth units during the experiment.

#### Water content determination

Samples were weighed immediately (fresh weight, FW), frozen in liquid nitrogen and stored at -80 °C before freeze-drying. After freeze-drying, the samples were weighed again (dry weight, DW). Water content (WC) was calculated as: (FW − DW)/DW.

#### Carbohydrate analysis

15-30 mg of ground sample powder was mixed in 1 mL ethanol 80% solution with mannitol (5 g. l^−1^) as internal standard. Non-structural carbohydrates (soluble sugar + starch content were determined according to Baffoin et al., 2021. High-performance liquid chromatography allows us to determine glucose, fructose, sucrose, and myo-inostitol. Starch was determined with classical enzymatic and spectrophotometric method.

#### Cytological analysis

For cytological analysis, 2020 growth units (n, Fig. 2) were cut with a razor blade and stored in fixing solution (3.7 % v/v formaldehyde, 5 % v/v acetic acid, 50 % v/v ethanol). The method described in Azri et al. (2009) was used to dehydrate and include the sample in London Resin White to increase the stiffness of the sample during the preparation of the cross sections. OmU2 rotary microtome (Reichert, Vienna, Austria) equipped with diamond knife was used to slice 4 μm thick cross sections. The sample was subsequently stained with 1 % toluidine blue/water (% m/v) during 30 s.

Each cross section was photographed on a Zeiss Axio Observer Z1 microscope equipped with a digital camera using the Zen imaging software system (Zeiss, Jena, Germany). The number of cells in cambium and xylem tissues was counted using 100x and 40x magnifications, respectively. Fiji software (Schindelin *et al*., 2012) was used for image analysis and counting was performed on 5 cell file by image.

### Modelling and Statistical analysis

#### Processing and modelling growth data

For primary growth, each growth curve was normalised by its maximal increment, the normalised growth was set to zero for the date before bud break (BBCH07) and then fitted using the classical Gompertz model (GM) (Rossi *et al*., 2003):

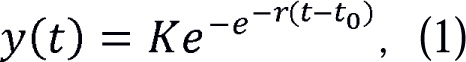

where *y(t)* is the fitted normalised growth (unitless), t is the time in days (day of year, DOY), K is the upper asymptote (unitless), t_0_ is the inflection point, i.e. time of maximum growth rate, and r is the normalised rate of change (in day^-1^).

The normalised growth rate (in day^-1^) is defined as the derivative *y’(t)* of the normalised radial growth with respect to time. For the Gompertz model, it is given as

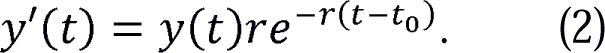

The date of growth cessation was defined as the day 95% of the total growth is reached (Fig. 3A).

**Figure 3:**
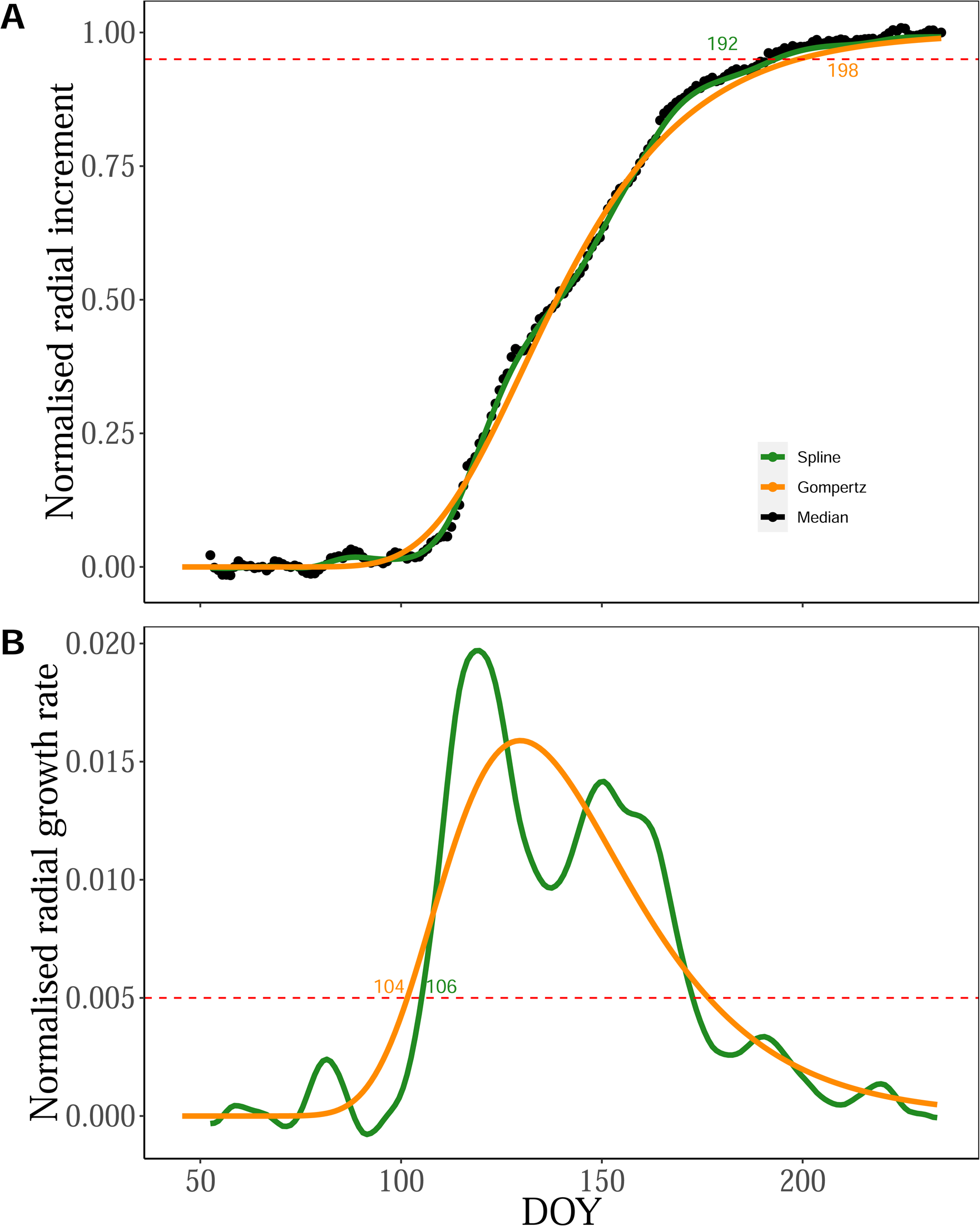
Example of the fit of the two models on the normalised values of the daily median for radial growth (A) and the growth rate i.e. the derivative of the two models. The red dotted lines represent the selection threshold for growth cessation (A) and growth onset (B).

Similarly each radial growth curve was normalised by its maximal radial increment.

Because the amount of data is much higher for radial growth, we could complement the use of the classical Gompertz model (GM) by a non-parametric model the cubic smoothing spline (de Boor, 1978). Cubic smoothing spline fits piecewise cubic polynomials to the points using a functional that balances attraction to the experimental points (Root Mean Square of the distance between all the points and their vertical projection on the curve) with a penalty on the wiggliness of the curve (RMS of the curvature). Splines offer more flexibility than the Generalised Additive Models (GM), and are agnostic (with no hypothesis on the shape of the growth curve) but require much more data to train.

For the GM on radial growth the equations are the same as (1) end (2), in which y is now the normalised radial growth rate and r the relative growth rate for secondary (Rossi *et al*., 2003): For the cubic smoothing spline we used the CSAPS Python library (https://github.com/espdev/csaps) to compute the splines (SpM). To avoid boundary effects, both sides of the raw curves were padded before spline computation with repeated values chosen as the average, respectively, over the first 10 days and the last 10 days. The splines were then analytically differentiated to get the normalised radial growth rates.

The dates of secondary growth onset and cessation were determined based on identical criteria for both models. The date of growth onset was defined as the first day the normalised radial growth rate exceeds a threshold value set at 0.005 day^-1^, with a minimal start date set at DOY 90 (Fig. 3B). The date of growth cessation was defined as the day 95% of the total radial growth is reached (Fig. 3A).

The counts for the number of cells in cambium and xylem tissues were fitted with a GM model (but not to a spline, due to the scarcity of the data). A value of two new cells produced was used for the onset of cambial activity. Consistent with the previous definition of growth in length and radius, cell production cessation was defined as the day when 95% of the final number is reached.

#### Statistical analysis

Statistical analyses were performed using R 4.0.5 (R Development Core Team) open-source software. Our experimental design is a cross-over experiment, each tree was his own control with at least one pair of branches, warmed and control. The effect of time and warming factor was tested by two-way ANOVA. Heteroscedasticity and variance homogeneity were assumed from residue distribution. Multiple comparisons were performed by comparing treatments at each time using the emmeans package (Lenth 2016). The relationships between physiological variables (water content, sugar content, and cell count) were investigated using Spearman correlation tests, as the data exhibited non-linear patterns with monotonic relationships.

## Results

### Warming branches advanced budbreak, but did not affect primary growth rates

The warming treatment had a significant effect on bud phenology, shifting the date of budbreak earlier in warmed branches compared with the control. The camera method and the visual observations provided convergent results regarding this effect of differential warming at the onset of primary growth (Fig. 4). Using the camera method, the date of 50 % increase in bud area was DOY 77 ± 5 and 83 ± 5 (mean ± SE) for warmed and control, respectively. With visual observation, budbreak (stage 07) occurred on DOY 80 ± 6 (mean ± SE) for the warmed branches and 14 days later (DOY 94± 9, mean ± SE) for the control treatment (Fig. 4). The camera method provided an earlier date for both treatments compared with the visual observations, and a smaller difference (6 days vs 14 days). For further bud developmental stages (i.e. 09 and 10 BBCH scale), the delay between both treatments remained similar with a difference of 14 days (Fig. 4). The shift was maintained during spring growth (Fig. 6A).

**Figure 4:**
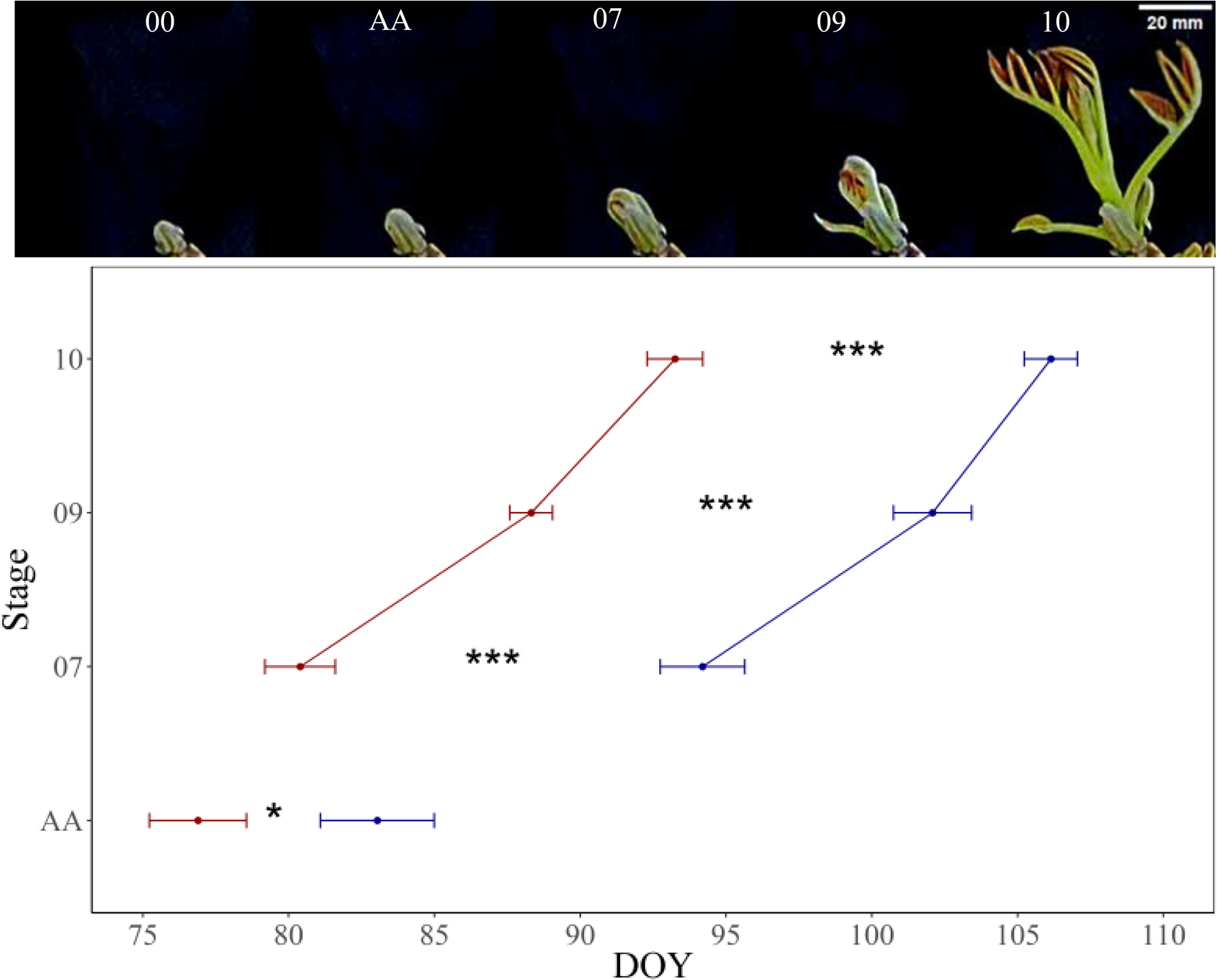
Phenological stage completion characterised the primary growth resumption of control and warmed walnut branches. Automated analysis (AA) allowed to determine the onset of budbreak based on a 50% increment in area (n= 10). Phenological stages were determined visually according to BBCH scale for each terminal bud (n= 50). Symbols and bars represent mean ± SE. The statistical significance of mean comparison is indicated (*<0.05 ; ** < 0.01 ; *** < 0.001) using pairwise comparison.

**Figure 6:**
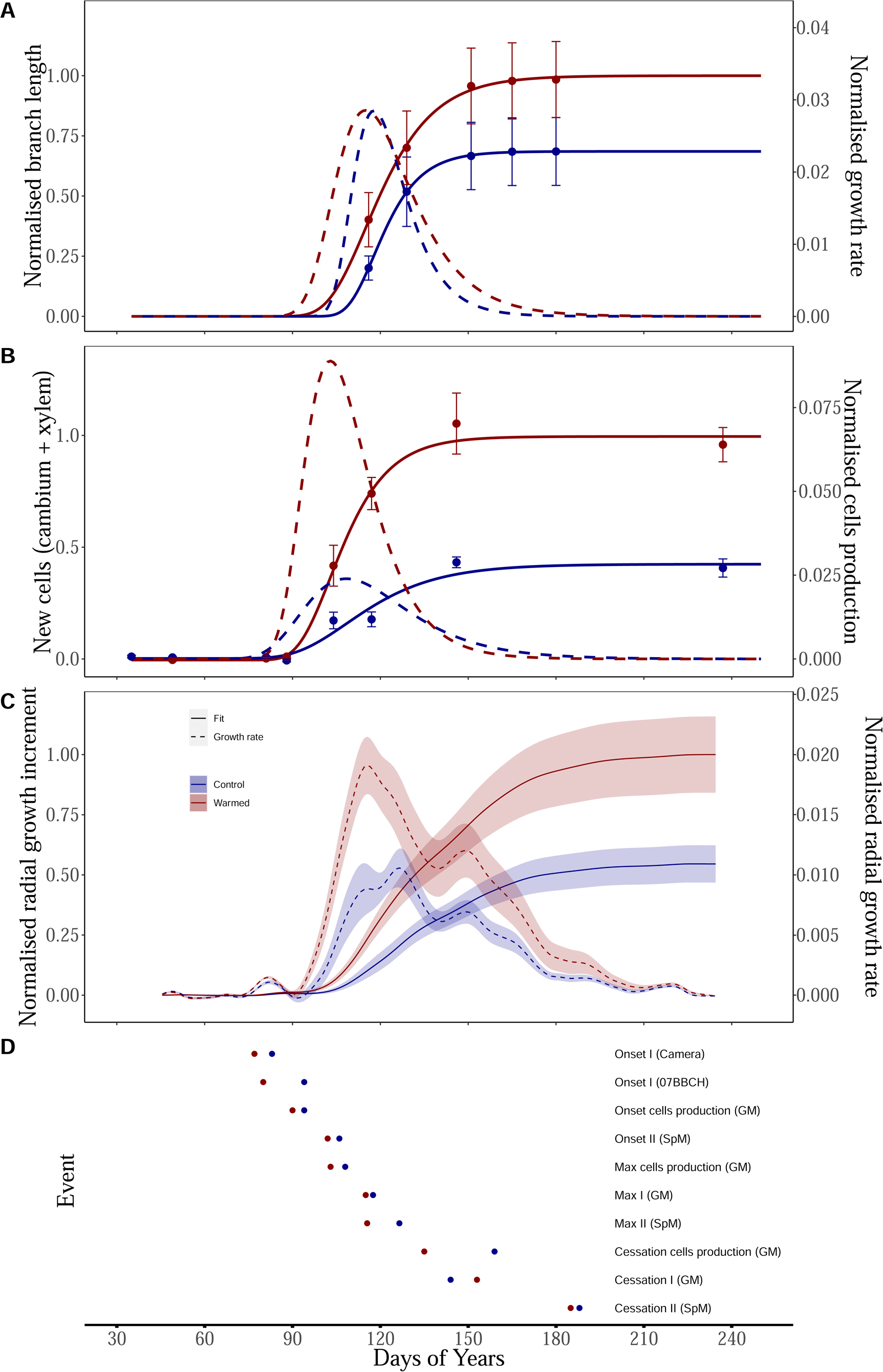
Time course of normalised (unitless) branch length (A), new cells (B), radial growth increment (C) and principal events (D) for warmed and control branches of walnut trees. Average fit with spline model (C) or Gompertz model (A&B) (solid line) and first derivative (dashed line). Thick line and error bar or colored area represent mean ± SE. For each variables, curves were normalised by asymptotic values of warmed treatment. Control branches and warmed branches are represented in blue and red, respectively. For each event, the mean date or the single value given by the model (in parentheses) is plotted.

At the end of the experiment, the length of the warmed branches was significantly higher than the control ones (Kruskall-Wallis test), by + 43 %. According to the Gompertz model (GM), warmed branches achieved 95% of length growth at 153 DOY, whereas control branches achieved it at 144 DOY (Fig. 6A&D). Taking into account the budbreak delay, the primary growth period of the warmed branches was 24 days longer than that of the control branches.

### Warming branches did not change secondary growth resumption, but increased growth rates and total wood cell production

The number of cambial cells in a radial file was stable around 5 until the end of March (Fig.5 A&F). A 1.6 fold increase was observed in April with a maximum value of 8 at DOY 104 and 114 for warmed and control branches, respectively, then a 2.7 fold decrease from 8 to 3 cells during August for both conditions (Fig.5 B, C&F). Concomitantly, the formation of new xylem cells started in April until the end of May (DOY 150), with twice more cells produced in warmed compared with control branches (Fig.5 D,E&G). According to GM, in both conditions, the onset of cell production, defined by the emergence of two new cells, occurred at DOY 90 for warmed branches and at DOY 94 for control branches. Cell production ceased (accounting for 95% of total production) at DOY 135 for warmed branches and DOY 159 for control branches (Fig. 5 B & D). However, these fitting approaches on non-paired and discontinuous sampling do not allow us to draw statistical conclusions regarding observed differences. Continuous diameter measurements, on the other hand, make this possible but results slightly differ depending on the model used, Gompertz Model (GM) or the Spline Model (SpM).

**Figure 5:**
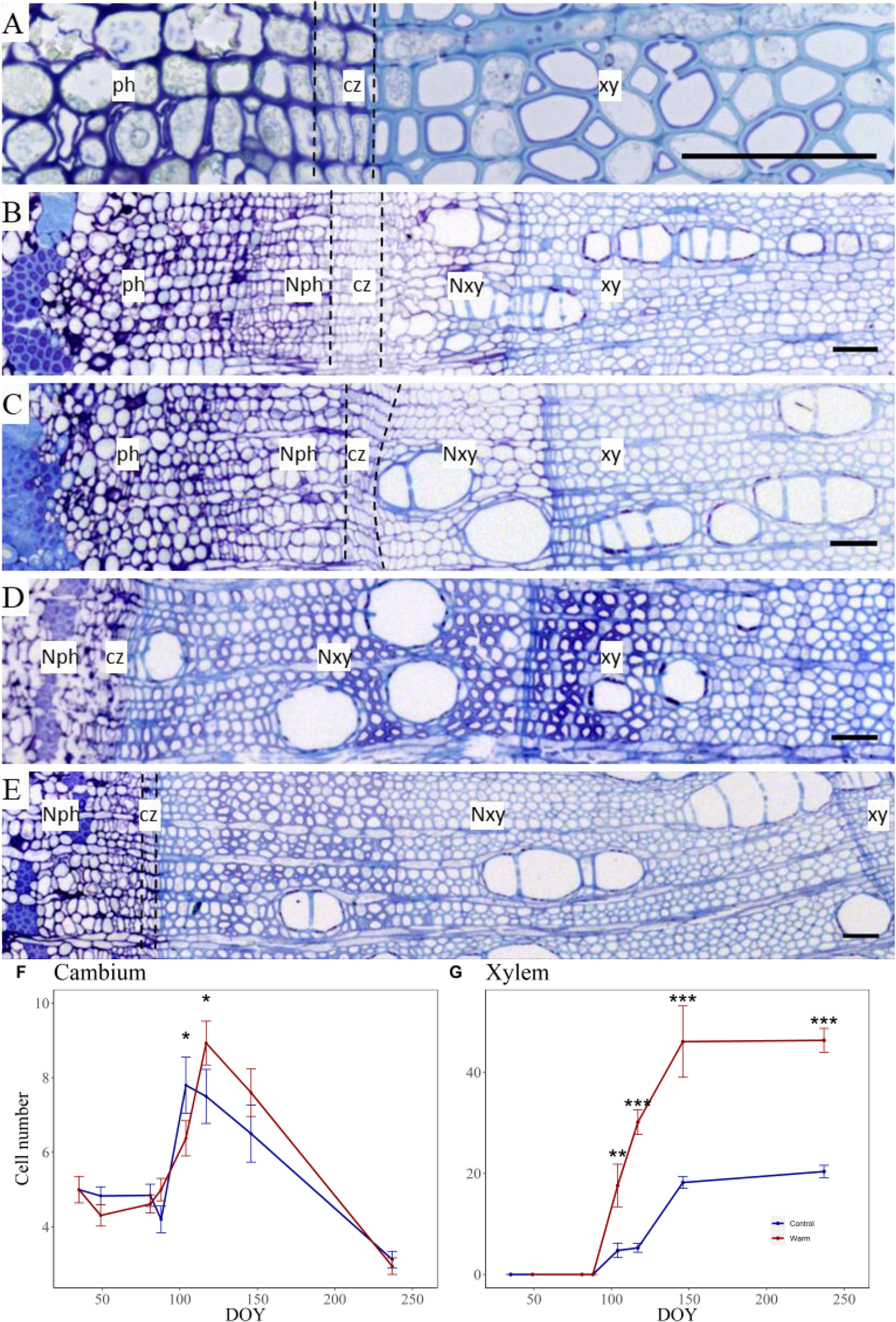
Transverse cross-section of the cambium, differentiating phloem and xylem, and mature xylem that were collected from control and warmed branches of Juglans nigra cv. Franquette. Dormant cambium was observed from 34 until 88 DOY for both treatments (A). Active cambial cell divisions were observed in control (B) and warmed branches (C) at DOY 104 (mid April). At the end of August (DOY 236), for control (D) and warmed branches (E), new cell formation stopped. cz, cambial zone; Ph, phloem; Xy, xylem; NPh, new phloem and NXy, new xylem formed during the experiment. Bars = 50 µm. Number of cambial cells (F) and new xylem cells (G) determined on transverse cross section during the whole experiment. Control branches in blue and warm branches in red. Mean ± SE, n= 5. The statistical significance of mean comparison is indicated (* <0.05 ; ** < 0.01 ; *** < 0.001).

When comparing the results between GM and SpM, a notable difference emerged. GM misfitted the data at the beginning and underestimated the start of secondary growth, 4 days earlier than the SpM (Table 1, Fig. 3 and Supplementary Fig. 2).

**Table 1:**
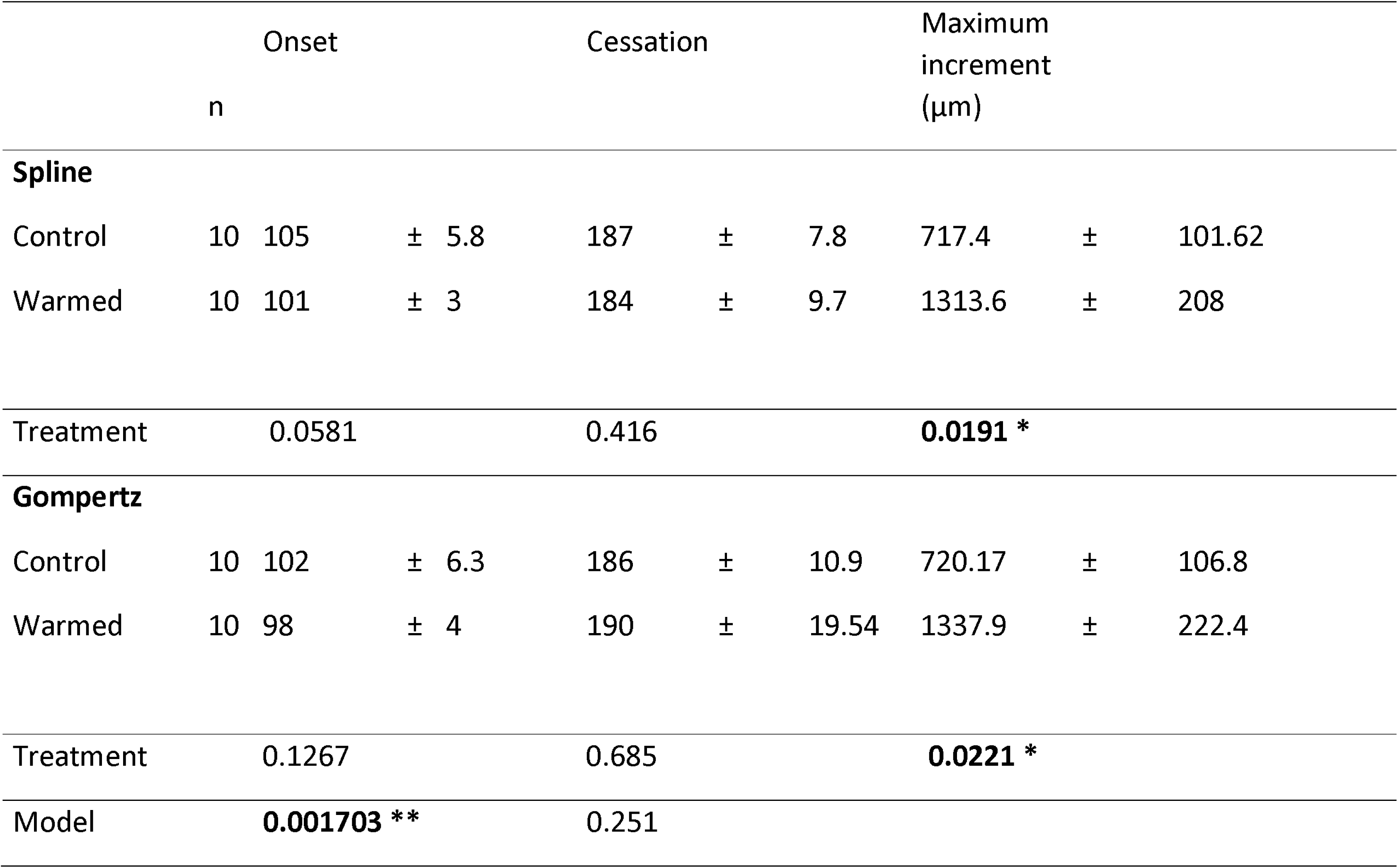
Secondary growth: Date of onset, cessation and maximum increment fitted by spline and Gompertz models for control and warmed branches (mean±SE). The date of growth onset was defined as the first day the normalised radial growth rate exceeds a threshold value set at 0.005 day-1, with a minimal start date set at DOY 90. The date of growth cessation was defined as the day 95% of the total radial growth is reached. One way Anova p-value were shown (* <0.05 ; ** < 0.01 ; *** < 0.001).

No matter the model, cessation of growth occurred at the same time for both conditions (insignificant difference) and for the same maximum radial increment values (Supplementary Fig. S2 and Table 1). Warmed branches were 1.8-fold larger than control branches (Table 1). According to GM, the inflection point, t_0_, and the normalised rate of change, r, were not found to be significantly different between the two conditions (Supplementary Table S1). The date of maximum growth rate (t_0_) occurred at 128 DOY for control branches against 125 DOY for warmed ones. In contrast, SpM displayed a bimodal shape in the growth rate curve, indicating two distinct peaks. The maximum growth rate did not occur at the same time for both conditions. Warmed branches reached their peak at DOY 116.5, while control branches reached theirs at DOY 127.5 (Fig. 6C&D).

To sum up the chronology of events (Fig. 6D), warmed branches initiated their primary growth earlier than control branches (DOY 77 vs. 83). Following budburst, cambial division first occurred around DOY 90 for both conditions, and the maximum division rate estimated by GM corresponds to the onset of radial growth increase (between DOY 102 and 108). Branch elongation and cell division ceased around day 150 of the year, with no strong certainty regarding any temperature-related differences. Secondary growth concluded simultaneously without temperature effects (DOY 185). Therefore, warmed branches displayed a longer primary growth and greater secondary activity.

### Warming promotes spring growth by early remobilisation of water and carbon

#### Warming modified water content dynamics for early primary growth

Water content (WC) exhibited disparities between the n and n-2 growth units. In the n growth unit, the WC of buds remained stable towards the end of winter (DOY 34 to 62), hovering around 0.65 g H_2_O.g^-1^ DW (Fig. 7A). Subsequently, it exhibited an increase during budbreak, occurring earlier for the warmed branches (DOY 81) in comparison with the control branches (DOY 88), although this difference lacked statistical significance. Soon after budbreak, the WC of newly formed branches reached its peak, surpassing 5 g H_2_O.g^-1^ DW (DOY 114). Following this peak, WC gradually declined throughout the remainder of the experiment until reaching a value of 1.16 g H_2_O.g^-1^ DW (average for both conditions). In n-2 growth units, WC demonstrated no differences between the two conditions (Fig. 7A). Unclear trends in rehydration could be observed from DOY 62 to DOY 114, ranging from 0.79 to 1.54 g H_2_O.g^-1^ DW. Then it dropped to 0.8 g H_2_O.g^-1^ DW until the end.

**Figure 7:**
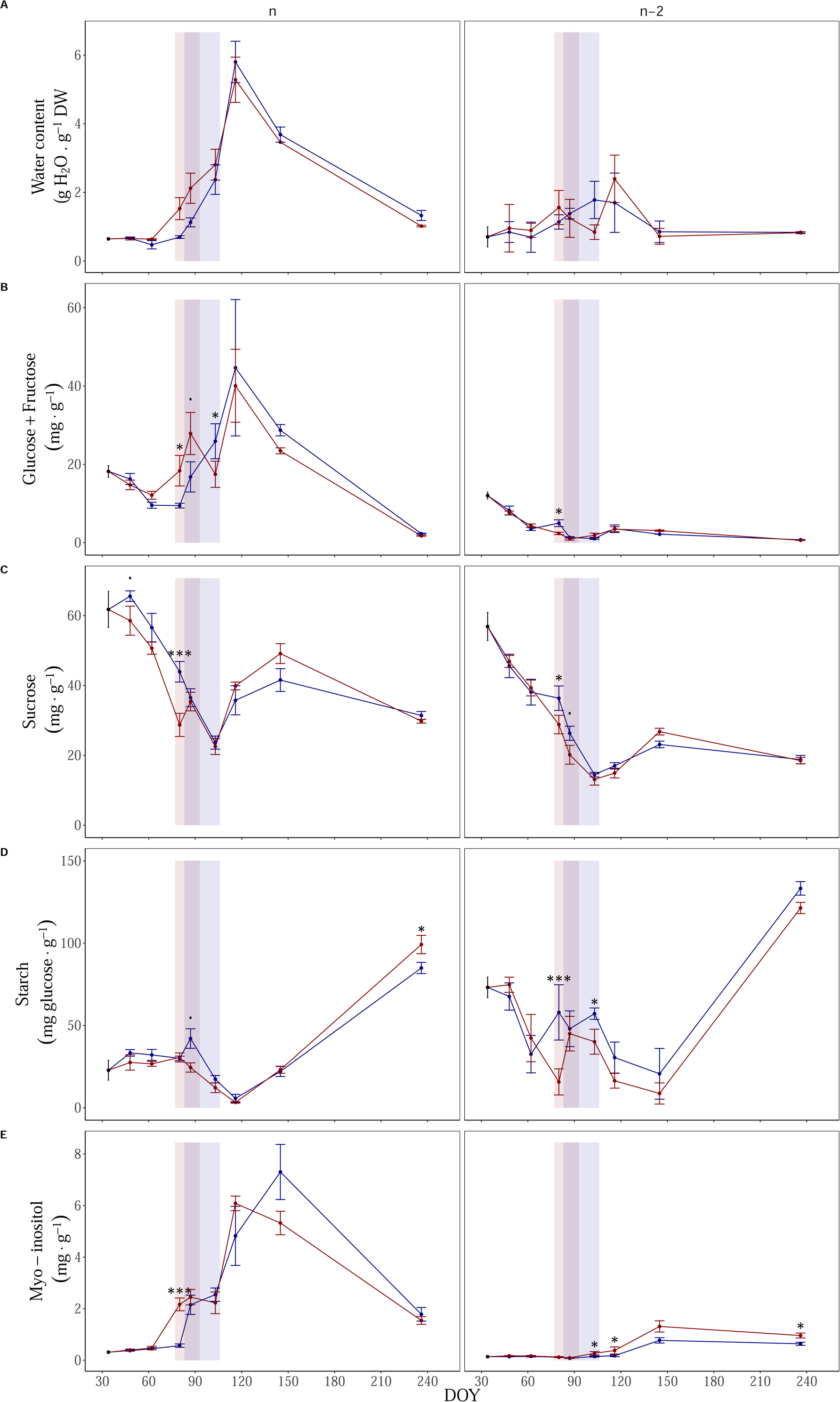
Seasonal course of of water content (A), major Solubles Carbohydrates (SC): glucose+fructose (B) and sucrose (C), but also starch (D) and Myo-inositol (E) for 2021 bud primary growth (n) and 2019 growth unit (n-2). The red and blue vertical bands represent the budbreak period between the dates obtained by the camera and the BBCH10 stage. Control branches in blue and warmed branches in red. Mean ± se, n= 5. The statistical significance of mean comparison by sampling date is indicated (* <0.05 ; ** < 0.01 ; *** < 0.001).

#### Warming modify bud carbohydrate remobilisation at the end of ecodormancy

For both treatments, the Glucose+Fructose (GF) content slightly decreased during the end of ecodormancy until the beginning of budbreak from 18.2 to 12.1 mg.g^-1^DW (DOY 62) for warmed and to 9.4 mg.g^-1^DW (DOY 80) for control branches (Fig. 7B). Then GF increased during the bud outgrowth phase. In warmed branches, the increase was observed earlier (DOY 88). Maximum GF contents (42 mg.g^-1^DW average) were observed for both conditions in mid-April during leaf expansion (DOY 114). In n-2 growth units, GF content was lower than in n growth units (12 mg.g^-1^ DW at the beginning) and had different dynamics. Indeed, there was a decrease by 3.5 in the GF content during the winter until April (DOY 104).

Inversely, during the same period, sucrose content in bud (n part) decreased significantly from 63 to 37 mg.g^-1^DW (Fig. 7C). Sucrose content tended to be slightly lower in warmed branches, with significative effects found at 49 and 88 DOY. Same trends were observed in n-2 parts, where sucrose content decreased from 57 to 23 mg.g^-1^DW and stayed slightly higher at DOY 80 and DOY 88 in control branches. Starch content was not impacted before bud break. Starch content was stable from February until March (Fig. 7D) then it decreased to a minimum (4.48 mg.g-1 DW average) observed at DOY 114. A starch peak was observed for control buds at DOY 88 but not for warmed branches. In n-2 parts, a decrease was observed sooner in March (DOY 62) for both treatments. Some discrepancies were noted during the budbreak, wherein warmed branches exhibited a diminished starch content in comparison to the control ones, marking a reduction of 75% on DOY 81 and 25% on DOY 104 (Fig. 7D).

#### Thermal differences fade after leaf out during active growth and starch accumulation

From April (DOY 114) until the end of the experiment in August (DOY 236), no difference appeared due to thermal treatment on GF content. For both conditions, it decreased from 42 to 2 mg.g^-1^ DW in newly formed branches (n part) and from 3.5 to 0.7 mg.g^-1^DW in n-2 growth unit (Fig. 7B). Conversely in n and n-2 part, sucrose doubled between DOY 114 and 145 (Fig. 7C). Then decreased at DOY 236, sucrose was 30 mg.g^-1^ and 18 mg.g^-1^DW, in n and n-2 part respectively. Over the same period, newly formed branches (n) exhibited a noteworthy increase in starch accumulation, statistically significant between conditions, 85 mg.g^-1^ DW for the control against 99 mg.g^-1^ DW for the warmed ones (Fig. 7D). In n-2 growth units, the starch content exhibited a large increase for both conditions, rising from May to August to reach a level as high as 125 mg.g^-1^, with no statistically significant difference between the two. Starch content tended to be lower in warmed branches during the secondary growth period.

#### Warming effect on primary and secondary growth modification reflected on Myo-inositol

There was a 4-fold increase of myo-inositol in the buds and growth unit of the year 2021 (n), observed on DOY 80 for the warmed treatment and on DOY 87 for the control (Fig. 7E). Contents stayed stable at least until the end of budbreak around DOY 104, then there was another significant increase to 6.08 mg.g^-1^ DW (DOY 114) for the warmed treatment and to 7.3 mg.g^-1^ DW (DOY 145) for the control, but without statistical difference between treatments. At the end of the experiment myo-inositol content dropped to an average of 1.66 mg.g^-1^ DW. For the n growth unit, myo-inositol was strongly correlated with water content (ρ =0.8, Fig. 8). Both were correlated to the number of cells in the cambial zone (ρ = 0.79 and 0.55 for myo-inositol and water contents, respectively). Secondly, in n-2 growth unit, there was an increase of myo-inositol content from April to the end of May from 0.14 mg.g^-1^ DW for both to 1.31 mg.g^-1^ DW for the warmed branches against 0.77 mg.g^-1^ DW for the control ones. At the end of the experiment (DOY 236), there was a significant difference between treatments: 0.96 mg.g^-1^DW for warmed branches against 0.64 mg.g^-1^ DW for control ones. This observed pattern over time followed the number of new xylem cells (n-1, ρ =0.9, (Fig. 7E & 5G). In other words, increased in myo-inositol followed the increase of new xylem cells produced in warmed condition.

**Figure 8:**
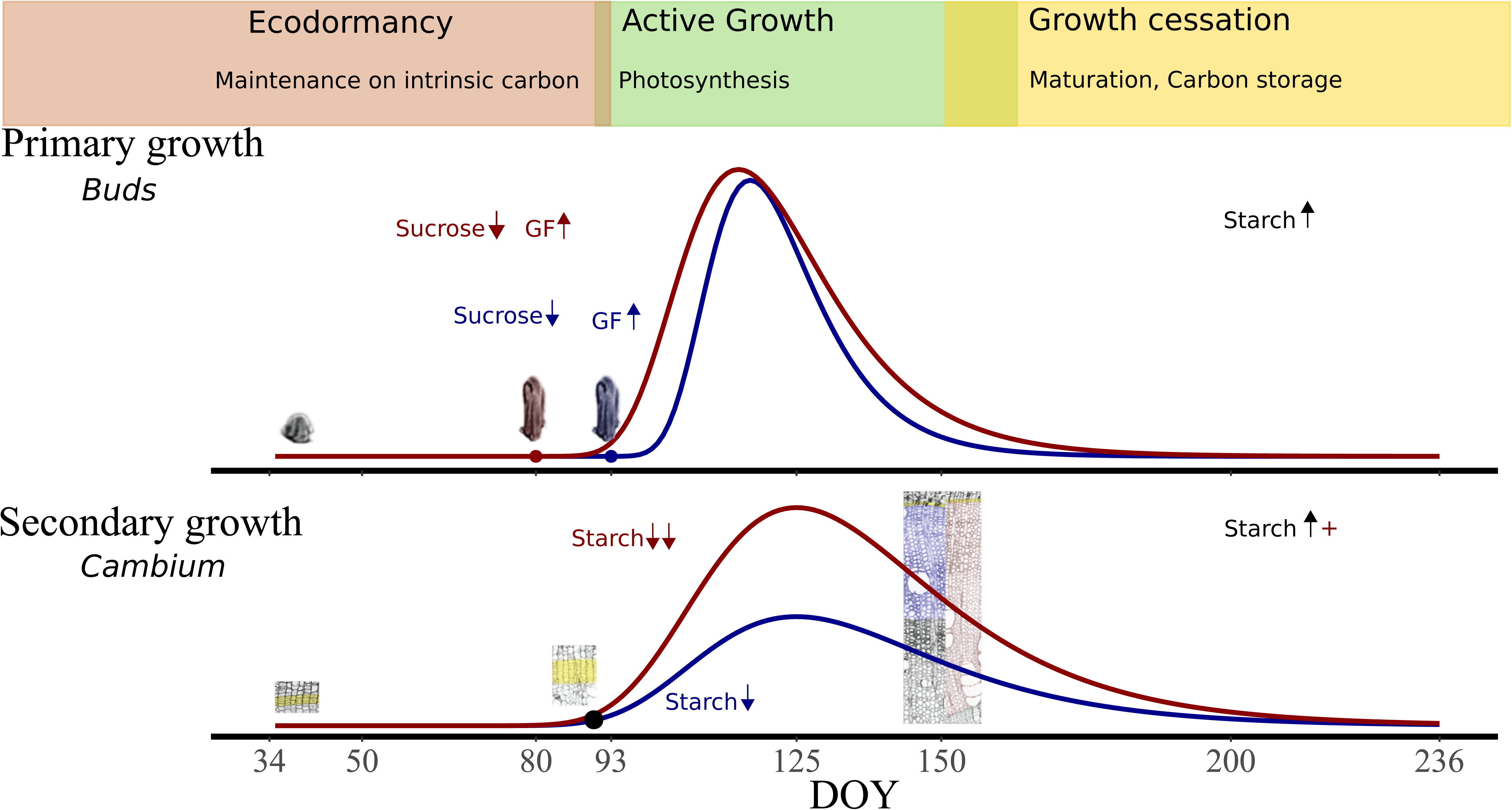
Summary of the temporal dynamics of primary and secondary growth rates between warmed and control branches. The hypothetical curves are directly adapted from the Gompertz model and excluded other environmental fluctuations. For buds, the transition from stage 00 to stage 07 (BBCH scale) is depicted. For cambium, the inactive, active, and end of cell production stages are shown; cambial cells are identified in yellow, and the new xylem in blue and red for control and warmed branches, respectively. Finally, the main differences in carbohydrate remobilisation, both in terms of timing and absolute content, are indicated. The up and down arrows illustrate an increase or decrease. Control in blue, warmed in red and black illustrates no difference.

## Discussion

### Warming advances primary growth onset, while secondary growth resumes independently to temperature but with higher cellular activity

Our study, from February to August, distinguishes three development phases: i) ecodormancy, characterised by plant maintenance and internal carbon consumption in the absence of leaves; ii) an active growth phase after bud break and cambial activity resumption; iii) a cessation phase involving the termination of branch elongation and cambium division before cessation of radial growth (Fig. 8).

#### Warming stimulates bud physiology during ecodormancy, hastening budbreak

Our main findings involve the boosting of budbreak due to temperature increase, regardless of the rest of the crown. According to the chosen thresholds and stages, in our conditions for walnut trees, budbreak defined as the start of swelling begins in mid-March, on DOY 77 for the warmed trees and DOY 83 for the control trees. Warmed buds were earlier and faster to budbreak. This is in line with prior researches. Targeted bud warming activated the growth of the bud independently of the rest of the tree (Bonhomme *et al*., 2013). Conversely, bark painting reduced branch temperature and delayed budbreak (Tixier *et al*., 2017*a*). Furthermore, experiments involving light/shading or change in albedo demonstrated that dormancy release is regulated at the bud level by the interplay of light and temperature (Zohner and Renner, 2015; Vitasse *et al*., 2021).

Since our trees were housed in late January, the bud endodormancy should have been released (23^rd^ January, inferred from the model from Charrier et al., 2018). Our warming experiment only plays on temperature and heat unit accumulation during ecodormancy stage. We observed early rehydration of the bud with earlier remobilisation of some soluble carbohydrates: glucose, fructose and myoinositol.

During ecodormancy phase and before budbreak, mobilised carbon (C) is intrinsic. Indeed, for both branches, carbohydrates were remobilised for maintenance respiration and metabolic costs associated to transfer and starch to soluble carbohydrates interconversions; their content thus decreased in bud and woody tissue until budbreak (Fig. 8). Starch storage becomes minimal during budbreak, at which point a sink-to-source transition likely occurred, and new carbohydrates were stored after leaf expansion, with maximum starch storage occurring at the end of the growing season. The carbon necessary for budbreak was likely sourced from the breakdown of branch starch, as previously indicated for *Quercus* and *Fagus* (Klein *et al*., 2016). However, in the case of walnut, evidence from literature using girdling and isotopic labelling experiments suggests that the C used for budbreak originates from more distant locations, such as the trunk, and its transport is required before budbreak for winter maintenance (Lacointe *et al*., 2004; Tixier *et al*., 2017*b*; Amico Roxas *et al*., 2021). In our experiment and at the bud scale, there is no significant effect of warming on carbon reserves. The availability of reserves is probably not a limiting factor for growth resumption. Reserves can be mobilised where needed without negative effects on control branches.

Temperature plays a crucial role in the rehydration of buds, leading to bud reconnection and the subsequent transport of water within trees. In the case of walnut trees at the end of winter, root pressure helps to restore hydraulic conductivity in the tree (Ameglio *et al*., 2002). Furthermore, soil temperature controls the date of stem rehydration but does not affect bud phenology (Charrier and Améglio, 2024). This restoration necessitates the presence of functional vessels, a characteristic that is typically not problematic for walnut trees, which are proficient at mitigating winter embolism (Améglio *et al*., 2001; Ewers *et al*., 2001). Moreover, increased root pressure is concomitant with budbreak (Cochard *et al*., 2001; Ameglio *et al*., 2002). There might be an interplay between xylem conductivity and bud rehydration and vascularisation. The hydration of the bud also depends on its reconnection to the vascular tissue through apoplastic or symplastic water transport. Furthermore, it is worth noting that temperatures thermodynamically influence the kinetics of metabolic reactions, potentially contributing to the overall process. Our results indicate that precise warming of branches leads to early rehydration and glucose-fructose remobilisation, along with early primary growth. The hydraulic reconnection between stem and bud appears to be a relevant direction to explore.

#### Asynchrony in meristems growth resumption

Our results indicate that bud and cambium resumptions were impacted in different ways by warming. Indeed, the resumption of cambial growth did not differ between warmed and control branches. By the end of March, on DOY 90, the cambium began to divide, while the bud was at stage 07 for the control branches and stage 09 for the warmed branches. The start of radial growth occurred in mid-April according to data from dendrometers and histology observation (Fig. 8). The shift in budbreak timing did not cause a shift in secondary growth resumption. This seems to indicate that primary and secondary growth resumptions are not simultaneously coordinated on walnut tree. According to these results, primary and secondary growth resumptions are slightly asynchronous. Indeed, budbreak occurs before stem increment, as for other diffuse porous like beech (Čufar *et al*., 2008; Kraus *et al*., 2016) or poplar (Li *et al*., 2013).

#### Warming enhanced cambial activity and wood production

After the resumption of meristem growth, the active growth phase begins. There was no difference in the primary growth rate between the warmed and control branches (Fig. 8). However the secondary growth rate was enhanced, probably through an increase in cellular activity at higher temperature. In other species, warming experiments have led to earlier cambial resumption (Begum *et al*., 2007; Gričar *et al*., 2007; Kudo *et al*., 2014). Oribe and Kubo in 1997 highlighted that localised warming modified evergreen coniferous (*Cryptomeria japonica*) cambium resumption but not deciduous ones (*Larix leptolepis*). However, asymmetric warming experiment on poplar revealed difference of secondary growth resumption between trunk faces (Begum *et al*., 2007). Conversely, *Quercus sessiliflora*, did not show any sensitivity to temperature in the timing nor the progression of wood and phloem formation. This observation was made on 150-year-old trees with much thicker and insulating bark than in other studies (Gričar, 2013). To explain these differences between species, it had been suggested that cambial rest is deeper and longer in deciduous than evergreen (Oribe and Kubo, 1997; Begum *et al*., 2007; Kudo *et al*., 2014). In 2008, Begum et al., proposed a period of several days above a threshold of 15°C daily maximum temperature before cambial reactivation for poplar. Our data do not establish any link between thermal time and cambial activity resumption (Supplementary Table S2). Additionally, the absence of differences in growth resumption dates between warmed and control samples does not allow us to propose a threshold for walnut.

The majority of studies examining the temperature effect induced by trunk or branch warming focus on growth resumption rather than the growth rate itself (Oribe and Kubo, 1997; Begum *et al*., 2007; Kudo *et al*., 2014). Nonetheless, a more extensive experiment conducted on Norway spruce by Gricar et al. (2007) revealed an increased rate of cell divisions at the beginning of the growing seasons in warmed trees. Our results revealed that walnut tree exhibits a higher cellular activity for warmed parts with higher cell division and radial growth rates during the active phase (Fig. 8).

Regarding sugars, it has been shown that non-structural sugars available in the cambial zone followed the intra-annual pattern of xylem formation, with a higher concentration when the growth rate was maximum (Deslauriers *et al*., 2009; Simard *et al*., 2013). However, our findings were less conclusive regarding water and sugar content determined in n-2 growth units. Notably, there were no discernible changes in water or carbohydrate content between the various treatments. The sampling part did not precisely target the cambium, hindering a comprehensive analysis of this aspect. Nonetheless, branches exhibited a tendency to rehydrate during primary growth and mobilise carbohydrate reserves. The cambium likely benefits from the concurrent remobilisation of stored carbon and water, occurring simultaneously with bud activation for growth resumption. Carbohydrate depletion stopped after the leaves appeared. This change probably indicates a shift from the local starch reserve, used as the main energy source for resuming growth, to the supply of C from photosynthesis after leaf fall. As a last point, Simard et al. (2013) highlighted that secondary growth limitation due to altitude was in response to sink activity controlled by temperature rather than the inability of trees to supply growing tissues with carbohydrates. Similarly to buds, carbon reserves are probably adequate to feed cambial growth resumption. The lack of pronounced differences between treatments does not allow for a conclusion regarding distinct sugar utilisation due to temperature asymmetry.

#### Coordination mechanisms between primary and secondary growth: is there an interplay between myo-inositol and auxin

Primary growth does not drive the initiation of wood formation. A disbudding experiment conducted by Kudo et al. (2014) revealed that cambial reactivation and vessel differentiation can occur independently of budbreak. This independent triggering, however, does not imply the absence of a connection between the two meristems. For example primary growth, potentially enhancing the supply of indoleacetic acid (IAA), seems to be indispensable for the continuous formation of wide vessel elements (Kudo *et al*., 2014). The model presented by Huang et al. in 2014 argues that primary growth and secondary growth are not synchronised at the beginning of wood formation but synchronisation occurs during the growing season. They emphasize that the coordination between primary and secondary growth processes throughout the growing season is crucial for optimal mechanisms to simultaneously allocate photosynthetic products and stored NSC for the growth of various tree organs. Along the same line, IAA has been characterised as young leaf signal, its polar transport from young shoot organs downward via the procambium and cambium induces and controls vascular differentiation (Aloni, 2013, 2015). Our experiment shows a significant correlation between the myo-inositol content in n part with the number of cambial cells and n-2 part content with new xylem cell formed. In the literature, myo-inositol content was already shown to be highly significantly correlated with the stem growth of *Pinus densiflora* trees (Kang *et al*., 2015). Other works pointed out that myo-inositol can form a conjugate with auxin (IAA) and so play a role in the storage and/or transport of this phytohormone (Kowalczyk and Bandurski, 1991; Loewus and Murthy, 2000). We could speculate that inositol might play an important role in transferring IAA from the bud to the cambium and adjacent xylem tissues. Moreover, the coordination of meristem through the interplay between carbohydrates and IAA seems relevant to be explored in future research.

#### Temperature unaffected growth cessation for primary and secondary meristems

Temperature did not affect the end of the primary nor secondary growth period. Indeed, around the 150^th^ day of the year, at the end of May or early June, the elongation of branches stopped, and no more new cambium cells were formed. Radial growth, measured by dendrometers, ceased around late July (DOY 180-190). Similar observations were made on warmed Norway spruce where continuously elevated temperatures did not significantly prolong cambial activity at the end of the vegetation period (Gričar *et al*., 2007). This observation indicates that internal and/or external factors (e.g photoperiod), should play a role in setting the end of meristem activity.

### Lack of compensatory mechanisms at crown scale could have a potential impact on tree architecture

The assumption that warming will cause earlier growth resumption in branches compared to control ones was partially confirmed at least for bud and this argues for some branch autonomy. Bud presents a certain autonomy as already shown with localised warming experiments (Bonhomme et al., 2010). This spring growth onset depends on carbohydrate reserve and allocation. It has been proposed in walnut that the bud is fed by sugar reserves in the trunk and that this transport involves the recirculation of water in the phloem and xylem according to Munch flow’s model (Lacointe *et al*., 2004; Tixier *et al*., 2017*b*). This remobilisation of reserve is associated with bud rehydration and frost dehardening (Charrier *et al*., 2013). Moreover, walnut is a diffuse porous wood with the ability of winter embolism resumption (Ameglio, 2002). The conductive tissues of walnut (n-1) are capable of transporting water and sugars (Lacointe *et al*., 2004). Consequently, primary meristem does not need new vascular tissues to deal with water and carbohydrates transport for primary growth. All those elements converge to explain an early warmed bud growth. Moreover, our results match the closest experiment in the literature on *Fraxinus lanuginose* (Osada et al., 2021). They characterised a shift in development between warmed and control branches with impact on leaf maturation on nitrogen allocation. It had also been theorised that branches had some autonomy regarding carbon allocation (Sprugel *et al*., 1991) but with some limits (Sprugel, 2002), for example, sink-to-source changes during the year. In 2004, Lacointe et al. conducted an experiment on walnuts subjected to asymmetrical lightning. Through the application of carbon labelling, they demonstrated branch-to-branch transfer, where newly sprouted shoots on each branch received a significantly greater amount of carbon mobilised from tree-wide reserves compared to reserves located locally on the mother branch. Their experiment also highlighted the remobilisation of reserves from the trunk to the branches, this transfer occurring during the winter. Even though, C is not likely to be limiting for primary growth resumption at tree level (Lacointe *et al*., 2004). This early demand for warmed branches could lead to an imbalance in distribution at tree crown level, resulting in competition for carbon resources.

Concerning cambium dynamics, even if warming does not alter growth resumption uniformly, the localised increase in activity suggests a degree of autonomy or, at the very least, a localised impact of warming. Numerous studies highlight a temperature-induced local effect confined to the treated region (Begum *et al*., 2007; Gričar, 2013). The augmented activity necessitates the utilisation of additional carbon reserves or photoassimilates for growth. This asymmetric response between two branches of the same tree could exert lasting effects on canopy architecture, whether through competition for reserves or by modifying the trade-off between early growth and frost hardiness. Although outside of these experimental conditions, other influential factors may come into play, temperature asymmetry effects could still influence the long-term development of the tree crown.

### Exploring the challenge of linking primary and secondary growth: emphasizing the significance of improved phenological analysis and methodology

Our work highlights the challenge of establishing a connection between primary and secondary growth for the concurrent determination of phenology. Primary and secondary phenological stage for onset and growth cessation should be well defined.

#### More accurate bud phenological assessment using a camera

As previously highlighted by Antunucci et al. (2015), budbreak has conventionally been associated with the final stages of bud development, specifically leaf unfolding (for example, Michelot et al., 2012; Takahashi et al., 2013). However, the initiation of primary growth actually started much earlier. We defined it when the bud began to swell due to rehydration. The traditional method for determining bud phenology is both time-consuming and with strong investigator effects (Liu *et al*., 2021). Using camera methods enable us to detect the resumption of bud growth earlier, with a difference of 3 to 10 days between treatments. The camera captured subtle changes, barely visible to the naked eye. This could serve as an effective proxy for bud rehydration. For our experiment, we employed phenoCam (automated digital time-lapse cameras) at the bud scale, although these cameras have previously been used at the tree level (Correia *et al*., 2020) or even at the landscape scale (Brown *et al*., 2016). We believe that the primary barriers for their application are their cost and the image processing time. Nevertheless, this approach is well-suited for remote sensing, and the processing algorithms are continually evolving (Correia *et al*., 2020).

#### Modelling radial growth: Gompertz vs spline

In this paper, we proposed a new and original way to analyse dendrometer data. Indeed, the Gompertz model (GM) approach had been widely used for xylem cell analysis and/or dendrometer data (Rossi *et al*., 2006; Michelot *et al*., 2012; Drew *et al*., 2022). GM has been chosen for good fitting performance on annual growth with easily interpretable parameters. GM is simply based on a sigmoidal growth hypothesis: slow growth that increases to a maximum and then decreases again. Regarding cytological analysis, GM was already outperformed by Generalised Additive Models (GAM) for describing intra-annual wood formation dynamics (Cuny *et al*., 2013). In our experiment, GM missed the correct date of growth resumption with a significant overestimation. The smoothing spline, just like GAM, is a non-parametric model which is more flexible than the parametric Gompertz model. It gave a better estimate of the onset growth and maximum growth rate. Moreover, the smoothing splines revealed a growth slowdown around may (140 DOY) due to a cold cloudy week (Figure S1 & S2). The Gompertz model could not detect such a transient slowdown. Our approach with dendrometer allowed us to infer cambial reactivation without destructive sampling and time consuming cytological and cell counting. However, analysis of the xylem cells indicated that there were more cells formed in the warmed branches and this information could only be obtained by cytological analysis, not via the dendrometers. More generally, cell counting and tracheidograms are more informative about the developmental processes than radial growth curves (Hartmann *et al*., 2017).

## Conclusion

We have demonstrated that temperature affects primary and secondary growth resumption differently. The discrepancy in growth resumption and activity between meristems were observed. Primary growth resumption was earlier but with no effect of temperature on the growth rate. This corresponds to an increase in the duration of the active phase. Secondary growth resumption was not modified by temperature; however, the growth rate was increased. The cessation of growth was not impacted by temperature and appeared to be similar between the two meristems. As a result, there was an asymmetry at the canopy level, with warmed branches being longer and thicker than the control ones. Although a difference in the remobilisation of carbon reserves was observed, reserves did not seem to explain the observed differences. Nevertheless, carbon signalling could play a role in coordinating the meristems during the active and cessation phases of growth. This specific point needs further exploration. Studying the signalling mechanisms, particularly the interplay between sugars and phytohormones, by which meristems perceive temperature, could provide new insights into meristem coordination. Examining the concurrent phenological dynamics in buds, terminal shoots, but also in trunk, and roots is imperative to understand their interconnection. This exploration could be carried out using predictive phenological models and tree function models. Furthermore, the influence of subtle spatiotemporal thermal variations on growth resumption could have a significant impact on tree architecture, especially in the event of late freeze events. Indeed, we could envision that early budbreak increases the risk of frost damage; late freezing events could alter architecture by killing early-warmed buds and causing a reorganisation of growth and carbon reserves at the crown scale.

## Acknowledgments

We express our gratitude to Maylis Urtebise for her valuable contributions during her internship on the experiment. Special thanks to Pascal Walser for his assistance in sampling, Brigitte Saint-Joanis for her skillful work in carbohydrates determination, Thierry Ameglio for some insights on dendrometer signal and Nicole Michac-Brunel for her expertise in cytological preparation. Additionally, we extend our appreciation to Marianne Lang for her artistic contributions in illustrating walnut branches for Figures 2 and 9.

## Author contributions

ND, BM, MS, FH, GC: Conceptualisation; ND, FH: Formal Analysis; BM: Funding Acquisition; ND, CS: Investigation; ND, CS, FH: Methodology; FH, GC: Project Administration; ND, FH: Validation; ND, FH: Visualisation; ND, FH: Writing – Original Draft Preparation; ND, FH, GC, BM, MS: Writing – Review & Editing

## Founding

This study was in part supported by multiple grant, “Contrat de Plan État Région Renoserre” to UMR PIAF, “Pack Ambition Recherche Doux Glace” and “Pari scientifique AgroEcoSystem INRAE”.

